# Nascent MSKIK peptide prevents or releases translation arrest in *Escherichia coli*

**DOI:** 10.1101/2022.10.03.510712

**Authors:** Teruyo Ojima-Kato, Yuma Nishikawa, Yuki Furukawa, Takaaki Kojima, Hideo Nakano

**Author notes:** Department of Agrobiological Resources, Faculty of Agriculture, Meijo University. Shiogamaguchi, Tempaku, Nagoya, 468-8502, Japan.

## Abstract

The insertion of the sequence encoding SKIK peptide adjacent to the M start codon of a difficult-to-express protein enhances protein production in *Escherichia coli*. In this report, we show that the increased production of the SKIK-tagged protein is not due to codon usage of the SKIK sequence. In addition, insertion of SKX, KKX, and AKX (X = G, L, H, Y, E, and F) at the N-terminus increased protein production. Furthermore, insertion of MSKIK just before the SecM arrest peptide (FSTPVWISQAQGIRAGP), which causes translational stalling on mRNA, greatly increased the production of the protein containing the SecM arrest peptide in the *E. coli* reconstituted cell-free protein synthesis system (PURE system). A similar phenomenon was observed for CmlA leader whose arrest is induced by chloramphenicol. These results suggest that the nascent MSKIK peptide prevents or releases translational stalling immediately following its generation during the translation process.

## INTRODUCTION

Despite protein production by microorganisms being an essential technology in life science research and industry, researchers still often encounter low productivity of target proteins in microbial hosts and mammalian cells such as Chinese hamster ovary cells (Kondo and Yumura, 2020; Krüger et al., 2022; Mathias et al., 2020). A general method to enhance the production of any protein in any organism without laborious trial and error will surely revitalize the bioeconomy that includes manufacturing bio-based products and bioenergy.

We reported previously that the insertion of codons 5’-TCT AAA ATA AAA-3’ or synonymous codons 5’-TCG AAG ATC AAG-3’ that encode four amino acids, Ser-Lys-Ile-Lys (SKIK), immediately after the Met start codon markedly increased the production level of proteins that were difficult to express in *E. coli* (Ojima-Kato et al., 2017a). Although we have shown that SKIK-tagging may promote translation but not transcription in *E. coli* in vivo expression experiments, the detailed mechanism remains unclear. Several recent studies showed that an optimal codon composition increases mRNA stability, resulting in improved protein production in *E. coli, S. cerevisiae*, and humans (Boël et al., 2016; Presnyak et al., 2015; Wu et al., 2019). Based on these observations, a possible hypothesis is that the codon composition encoding SKIK may influence protein production.

Newly generated polypeptides produced during translation are called nascent chains. These chains have important roles in various cellular processes including their maturation and quality control of proteins and mRNA, and can be involved in the control and regulation of cell-wide biological phenomena (Choi et al., 2018; Niwa et al., 2019). Among them, nascent chains that cause translational stalling are called ribosome arrest peptides (RAPs). RAPs contain specific amino acid sequences that interact with internal components of ribosomes in the nascent state. Various RAPs with different arresting mechanisms are distributed widely across species (Emmanuel et al., 2019; Matsuo et al., 2020; Phadtare et al., 2002; Yanagitani et al., 2011).

SecM, a bacterial secretion monitor protein composed of 170 amino acid residues, is a well-known RAP in *E. coli* (Ito and Chiba, 2013). Translational stalling by SecM is caused by the exit tunnel of the ribosome 50S subunit interacting with the nascent chain FXXXXWIXXXXGIRAGP (X is any amino acid) present in the C-terminal region of SecM (Nakatogawa and Ito, 2002; Zhang et al., 2015). This RAP functions as an in vivo or in vitro translation arrester in *E. coli* even when inserted into another protein of interest (Contreras-Martínez and DeLisa, 2007). Other types of RAPs such as the CmlA leader peptide (KNAD) induced by chloramphenicol, TnaC (WFNIDNKIVDHRP*, where star represents a stop codon) induced by Trp, and the artificially designed WPPP sequence (FQKYGIWPPP), are also profiled in *E. coli* (Ito and Chiba, 2013).

In this study, we demonstrate how SKIK-encoding codon usage affects protein production in *E. coli* in vivo and in vitro expression systems to verify whether the codons of SKIK or the amino acid sequence are important for translation enhancement. We then hypothesize that the SKIK peptide, the nascent chain, may act as a translation enhancer, preventing translational slowdown caused possibly by the interaction between the ribosome and nascent polypeptides. We also evaluate the effect of combinations and orders of SKIK or MSKIK with various RAPs. Finally, we reveal that nascent MSKIK can strongly prevent or release translational stalling that RAPs may cause.

## RESULTS

### SKIK encoding codons do not affect protein production in *E. coli* in vivo and in vitro expression systems

In our previous study, two patterns of SKIK encoding codons (5’-TCT AAA ATA AAA-3’ and 5’-TCG AAG ATC AAG-3’) were shown to affect protein synthesis to a similar extent (Ojima-Kato et al., 2017a). In this study, we evaluated all synonymous codons except the last lysine codon (AAA) and quantified the yield of His-tagged proteins by dot blot analysis using anti-His tag horse-radish peroxidase (HRP) and ImageJ software.

In isopropyl-β-D-thiogalactopyranoside (IPTG)-induced *E. coli* samples, the production of each SKIK-tagged rabbit scFv was approximately 20–28 times higher than that of no-SKIK tagged-scFv, and the 36 variants showed almost the same expression level regardless of the different codon usage (Figures 1 and S1). Analysis of variance showed no significant differences in protein production levels among these 36 samples. Thus, the results showed no correlation between the codon patterns encoding the SKIK peptide tag and the amount of the expressed protein.

**Figure 1.**
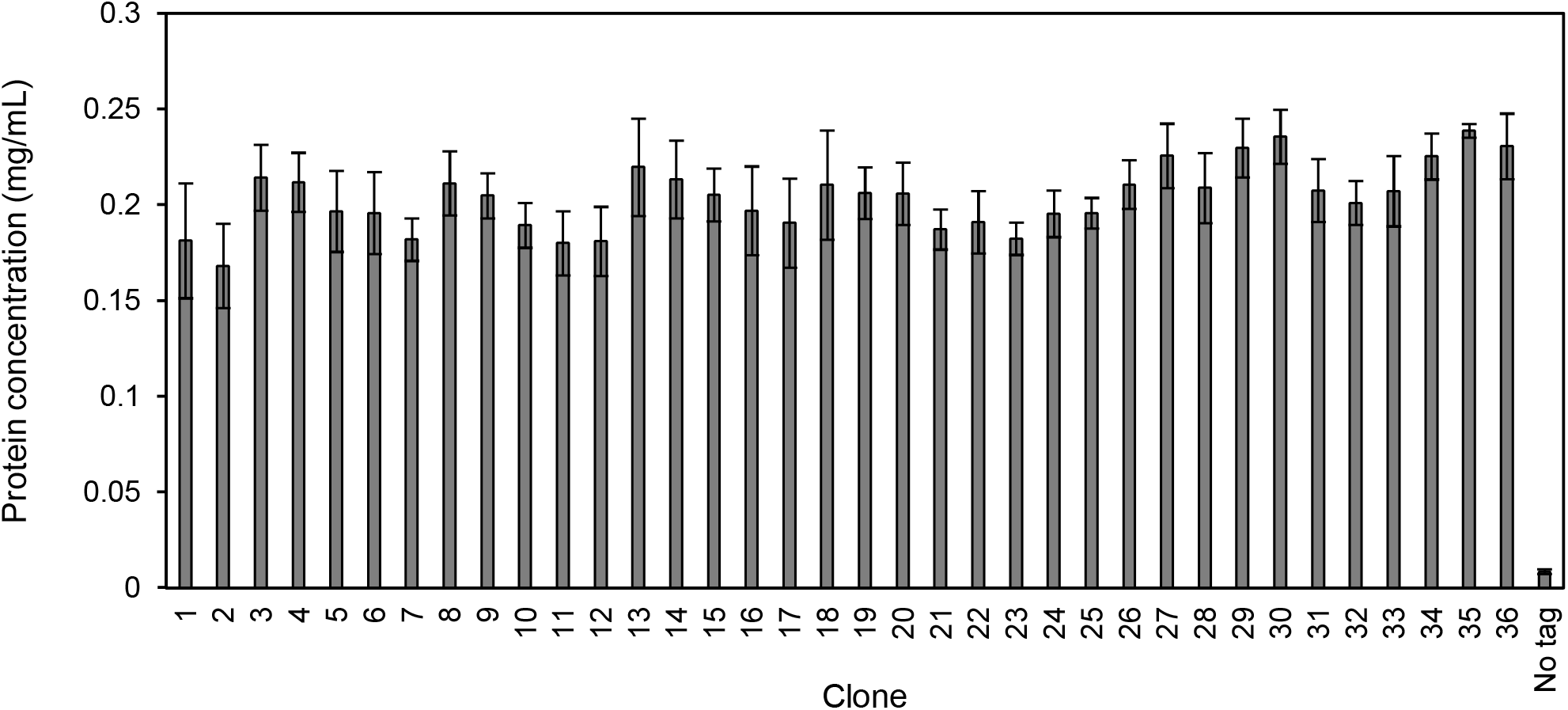
Protein production in *E. coli* of an antibody fragment tagged with SKIK composed of various codons. In vivo protein (single chain antibody fragment from rabbit fused with a C-terminal His-tag) expression in *E. coli* was performed by IPTG induction. The dot-blotted proteins in cell lysates were visualized and quantified with the HRP conjugated antibody against the His-tag to react 3,3’,5,5’-tetramethylbenzidine (TMB). Images were analyzed with ImageJ software. Sample numbers correspond to the SKIK-encoding codon patterns listed in Table 1.

The same 36 variants of SKIK-tagged scFv were expressed by auto-induction for 72 h in TB medium at 30°C. Protein productivity was analyzed by SDS-PAGE, followed by CBB staining, yielding clear bands of all SKIK-tagged scFvs, whereas no target band was observed without the SKIK tag (Figure S2). There was no significant difference in the amount of protein produced among the 36 variants. These results indicated that adding the SKIK tag increases the production of target proteins uniformly, and this increase is independent of codon usage or induction methods. Moreover, the in vitro *E. coli* reconstituted cell-free protein synthesis system yielded good expression at similar levels for the 36 variants, whereas the non-tagged protein was not detected (Figure S3). On the basis of the above results obtained for in vivo expression using different induction methods and in vitro expression, we concluded that the SKIK peptide tag increases the amount of protein produced both in vivo and in vitro in the *E. coli* expression system and is not affected by codon usage in either case.

**Table 1.** The codon patterns encoding the SKIK peptide tag.

| Clone No. | S | K | I | K |
| --- | --- | --- | --- | --- |
| 1 | tct | aaa | att | aaa |
| 2 | tct | aaa | atc | aaa |
| 3 | tct | aaa | ata | aaa |
| 4 | tct | aag | att | aaa |
| 5 | tct | aag | atc | aaa |
| 6 | tct | aag | ata | aaa |
| 7 | tcc | aaa | att | aaa |
| 8 | tcc | aaa | atc | aaa |
| 9 | tcc | aaa | ata | aaa |
| 10 | tcc | aag | att | aaa |
| 11 | tcc | aag | atc | aaa |
| 12 | tcc | aag | ata | aaa |
| 13 | tca | aaa | att | aaa |
| 14 | tca | aaa | atc | aaa |
| 15 | tca | aaa | ata | aaa |
| 16 | tca | aag | att | aaa |
| 17 | tca | aag | atc | aaa |
| 18 | tca | aag | ata | aaa |
| 19 | tcg | aaa | att | aaa |
| 20 | tcg | aaa | atc | aaa |
| 21 | tcg | aaa | ata | aaa |
| 22 | tcg | aag | att | aaa |
| 23 | tcg | aag | atc | aaa |
| 24 | tcg | aag | ata | aaa |
| 25 | agt | aaa | att | aaa |
| 26 | agt | aaa | atc | aaa |
| 27 | agt | aaa | ata | aaa |
| 28 | agt | aag | att | aaa |
| 29 | agt | aag | atc | aaa |
| 30 | agt | aag | ata | aaa |
| 31 | agc | aaa | att | aaa |
| 32 | agc | aaa | atc | aaa |
| 33 | agc | aaa | ata | aaa |
| 34 | agc | aag | att | aaa |
| 35 | agc | aag | atc | aaa |
| 36 | agc | aag | ata | aaa |

The minimum free energy of the thermodynamic ensemble from the entire transcripts (mRNAs from transcript start position GGG to T7 terminator) calculated by RNAfold was – 311.1 ± 12.6 kcal/mol for all SKIK tagged clones, whereas the value was –273.0 kcal/mol for the non-tagged clone. Specific nucleotides from the 5’ UTR (–19) TTAAGAAGGAGATATACAT to ACCCAGACTCCAGCCTCC in r1scFv were calculated to be –7.3 ± 1.0 kcal/mol in SKIK-tagged variants, and –6.5 kcal/mol in the no-tag variant (Figure S4).

### Increased production of the SKIK-tagged protein is caused by improving translation and not transcription

To evaluate whether the addition of the SKIK peptide tag affects transcription or translation, in vitro transcription and translation were performed separately using rabbit antibody genes L- and H-chains (GenBank accession numbers BAX56586 and BAX56587) (Ojima-Kato et al., 2017b), with and without SKIK tags. The amounts of mRNA transcribed from each template were similar when equal amounts of PCR products were used as templates, regardless of the presence or absence of the SKIK tag (Figure 2A). In contrast, adjusting the template mRNA amounts to be equivalent and performing the translation reaction in the PURE system showed an increase in protein production of both the H- and L-chains when the SKIK tag was present (Figure 2B). Thus, adding the SKIK tag for in vitro protein synthesis does not alter transcription but improves translation significantly.

**Figure 2.**
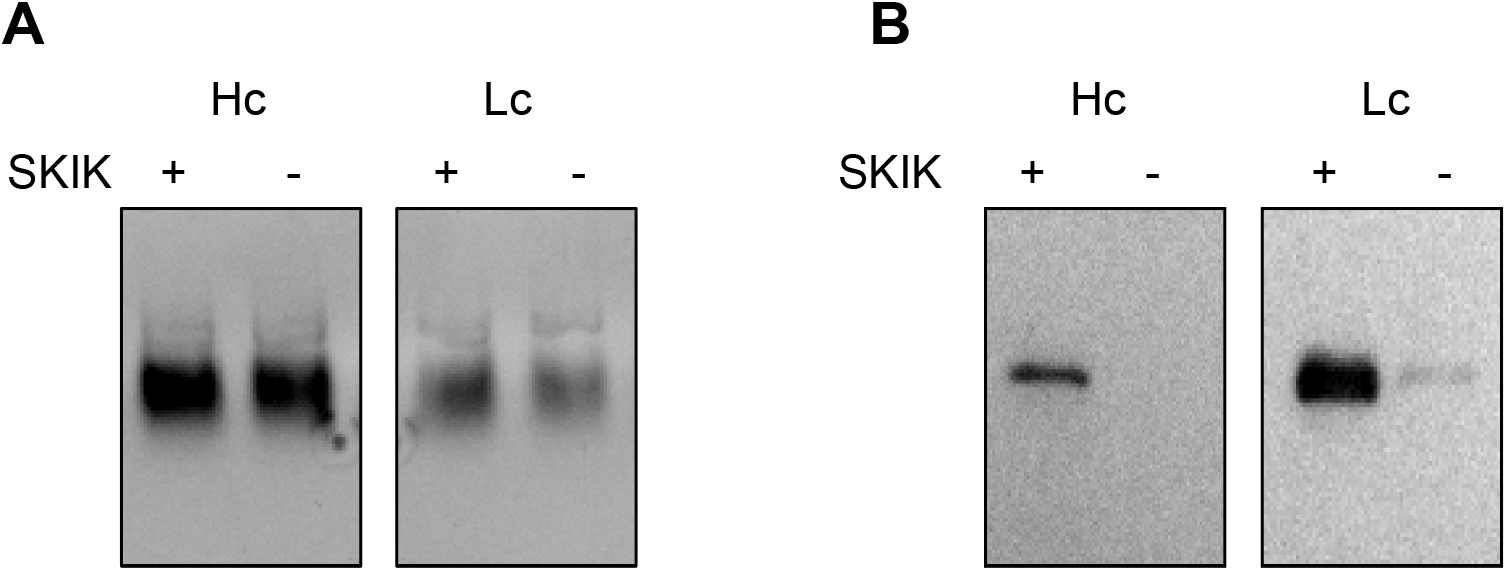
In vitro transcription and translation of rabbit antibody genes with/without the N-terminal SKIK peptide tag. Rabbit heavy chain (Hc) and light chain (Lc). Plus and minus indicate the presence and absence of the SKIK tag, respectively. (A) Transcript products (1 μL from 25 μL reaction volume) in agarose electrophoresis. The bands were visualized with ethidium bromide staining and UV light excitation. (B) Western blotting analysis of cell-free protein productions using mRNA as templates. The HA-tagged Hc and His-tagged Lc were detected with anti-HA and anti-His tag HRP-conjugated antibodies, respectively. The HRP reaction was carried out with TMB blotting solution.

### Peptide tags with only three amino acid residues SKX, KKX, and AKX can improve protein production

The above results indicate that the amino acid sequence of the SKIK tag is important for promoting translation. Therefore, the effect of other amino acids at the second residue position and adjacent to the starting M was examined because this position was reported to be important for protein production (Bivona et al., 2010; Boël et al., 2016). Here, the production of a rabbit scFv in *E. coli* was evaluated with a variety of tag sequences (1^st^: S, K, or A; 2^nd^: K; and 3^rd^: G [codon GGT], L [TTA], H [CAT], Y [TAT], E [GAA], or F [TTT]). Surprisingly, all of the N-terminal tagged-scFvs were more productive when compared with the scFvs without the N-terminal tag (Figure S5).

### MSKIK peptide tag counteracts ribosome arrest caused by SecM

The obtained results suggest that the amino acid sequence, but not the mRNA sequence, of the SKIK peptide tag contributes to translation enhancement. We then evaluated how combining a typical SecM arrest peptide (SecM AP, FSTPVWISQAQGIRAGP) and the SKIK peptide affected translation in an effort to understand the mechanism of the SKIK tag on translation. The amino acid sequences of the eight constructs are shown in Figure 3A. The PURE system was used for transcription and translation reactions, and the produced proteins were analyzed by Western blotting with the anti-His tag antibody conjugated with HRP. The amount of proteins containing SecM AP was below the detection limit (Figure 3B, lanes 1 and 5), whereas the protein without SecM AP was detected (Figure 3B, lanes 3 and 4). This observation implies translation arrest by SecM AP occurred in the PURE system, resulting in poor protein production.

**Figure 3.**
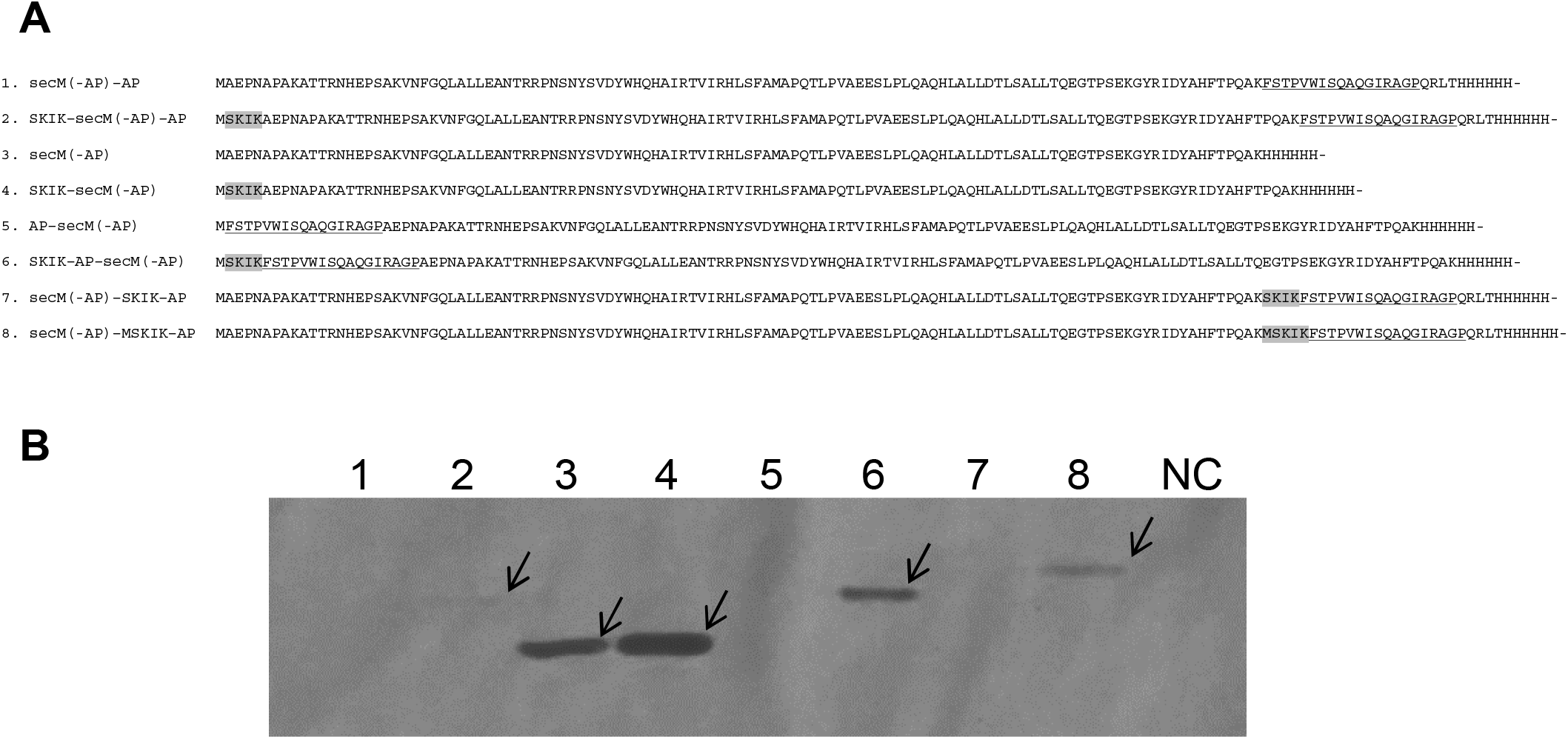
Tested constructs containing SecM AP and their Western blotting analysis. (A) Amino acid sequences of the eight constructs. Insertions of SKIK or MSKIK are shaded in grey. Parts of the SecM arrest peptide (FSTPVWISQAQGIRAGP, named AP) are underlined. The sequence AEKP to PQAK is common to all constructs, and SecM (-AP) is the non-arrest part of SecM. (B) Western blotting analysis after *E. coli* cell-free protein synthesis. The His tag at the C-terminus was detected. Arrows indicate the detected target bands. NC represents the negative control without the DNA template.

Surprisingly, N-terminal SKIK placed immediately before the SecM AP increased the amount of synthesized protein significantly (lane 6). This effect was not as noticeable when the distance between the SKIK tag and SecM AP was 112 amino acid residues (lanes 1 and 2). Interestingly, the insertion of the sequence MSKIK adjacent to SecM AP in the middle region of the protein gave higher production than the insertion of SKIK in a similar construct (lanes 7 and 8). This result implies that the presence of this tag not only at the N-terminus but also in the middle of ORFs, just before the AP, may have a translation-promoting effect on entire genes.

These results revealed that the SKIK tag containing the initiation codon encoding M placed just before the SecM AP releases or reduces ribosome arrest and promotes translation of the full-length gene. In addition, the presence of M and the physical distance from the AP are important factors that define the strength of the translation enhancement.

### MSKIK peptide prevents or releases ribosome arrest induced by chloramphenicol

Next, we assessed the effect of MSKIK on SecM AP and other types of APs, including CmlA leader whose arrest is induced by chloramphenicol and the artificial arrest sequence WPPP, by fusing the super folder green fluorescent protein (sfGFP) gene downstream of each AP (Figure 4A). Tagging with the SKIK peptide caused an increase in the fluorescence intensity of AP-fused sfGFP from 2.7 × 10^2^ to 2.4 × 10^3^ (8.9-fold) for SecM AP (FSTPVWISQAQGIRAGP) and 1.8 × 10^4^ to 4.3 × 10^4^ (2.4-fold) and 4.3 × 10^3^ to 9.3 × 10^3^ (2.2-fold) for CmlA leader (KNAD) in the presence of 0.1 and 1.0 μg/mL of chloramphenicol, respectively. In contrast, the fluorescence was weak for CmlA leader at a high concentration (10 μg/mL) of chloramphenicol without and with the SKIK tag, namely 4.3 × 10^2^ and 5.6 × 10^2^, respectively. Moreover, weak fluorescence was observed for WPPP (FQKWGIWPPP) with and without the SKIK tag with values of 4.6 × 10^2^ and 7.3 × 10^2^, respectively. This result suggests that ribosome arrest caused by CmlA leader and SecM were also reduced by MSKIK, which increased sfGFP expression. The same samples were also subjected to SDS-PAGE followed by Western blotting against the C-terminal His-tag (Figure 4B). The band intensities in this assay also correlated with GFP fluorescence intensities for all samples. This correlation shows that the fluorescence intensity of the sfGFP assay reflects the quantity, but not the quality, of the translated protein.

**Figure 4.**
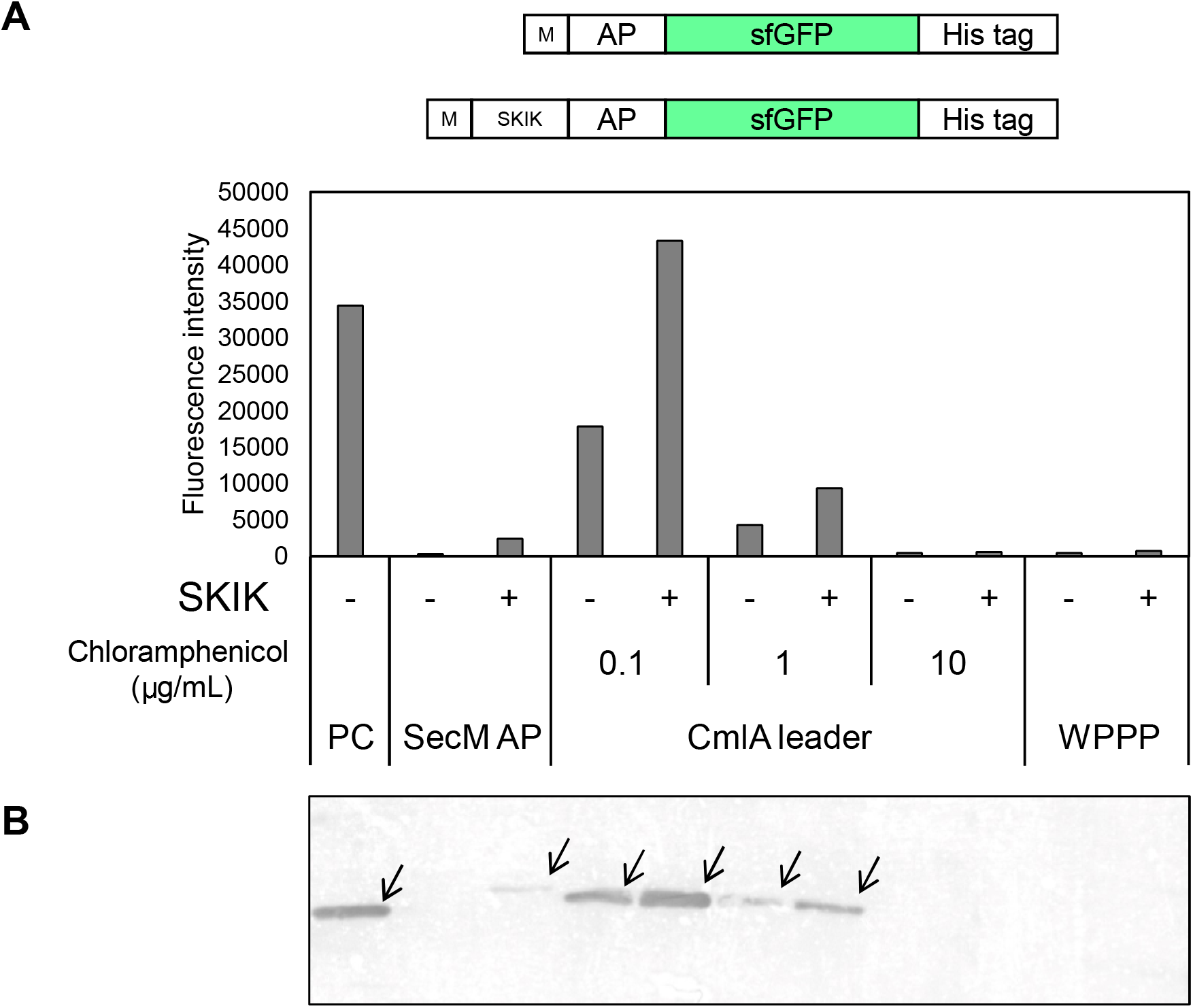
Effect of N-terminal SKIK peptide tag on various arrest sequences in cell free-protein synthesis. (A) Fluorescence intensity of sfGFP. The illustration above the graph depicts the genetic constructs compared. (B) Western blotting analysis. The His-tag at the C-terminus was detected. Arrows indicate the detected target bands. The order of samples is the same as that shown in A. +/-indicates the presence/absence of SKIK immediately upstream of each arrest. PC: positive control without arrest peptide; SecM AP: FSTPVWISQAQGIRAGP; CmlA leader: KNAD; and WPPP: FQKYGIWPPP.

## DISCUSSION

We showed that a short nascent chain, MSKIK, increases the production of difficult-to-express proteins by likely preventing or releasing ribosome arrest caused by RAPs. Various studies have demonstrated that the codon usage of a gene affects the amount of protein produced (Goodman et al., 2013). In particular, codon bias encoding the N-terminus affects mRNA expression levels but has been reported to be largely independent of mRNA translation or mRNA stability (Zhou et al., 2016). Boël et al. reported that the initial ∼16 codons more likely influence mRNA folding, rather than the entire codon content, by directly modulating translation efficiency and mRNA stability (Boël et al., 2016).

In the present study, we evaluated protein production in all synonymous codon patterns in SKIK except for the last K and showed that all codon patterns yielded the same protein production enhancement effect as observed for the original SKIK tag. In general, the presence of rare codons reduces protein expression (Liu, 2020). Interestingly, SKIK containing rare codons such as ATA (Ser) and TCG (Leu) (Sharp and Li, 1986) gave protein levels that were consistently increased. The energy and structure of the RNA were examined for each codon pattern, with no distinct differences found (Figure S4). This finding is also consistent with the observation that there was no significant difference at the protein level among the 36 variants.

In addition, SKX, KKX, and AKX immediately after the starting M increased protein production (Figure S5). This observation supports the hypothesis that this increased protein production is codon-independent. Because the second K was fixed and the third amino acid was evaluated by arbitrarily selecting representative amino acids with different properties, this limited evaluation is incomplete, but at least SK, AK, and KK, after the M may be important for enhancing protein production.

Separately conducted in vitro transcription and translation experiments revealed that the SKIK tag did not affect the amount of transcription product but only increased translation (Figure 2). In our previous *E. coli* in vivo experiment, the mRNA level quantified with real-time PCR and the amount of mRNA per cell decreased when using the SKIK tag, whereas the amount of protein increased by ∼30-fold (Ojima-Kato et al., 2017a, p.). Taken together, SKIK-tagging significantly increases the efficiency of *E. coli* in vivo and in vitro translation. However, because the present study only compares the amount of products, further detailed analysis is required to determine whether there is a change in the transcription/translation rate due to SKIK-tagging.

Combining SecM AP and SKIKs revealed unexpectedly that placing MSKIK immediately before the AP counteracts the effect of the arrest and increases protein production. In addition, comparative experiments with MSKIK and SKIK inside the ORF showed that the effect of MSKIK was stronger. This observation suggests that the nascent chain, MSKIK, at the N-terminus and inside the ORF cancels the arrest caused by SecM immediately after this nascent chain is generated. The kinetics of translation elongation are critical for driving gene expression and maintaining proteostasis, including nascent chain folding and binding to different ribosome-associated machinery (Stein and Frydman, 2019). The ribosome arrest mechanism of SecM has been analyzed extensively (Bhushan et al., 2011; Gumbart et al., 2012; Tsai et al., 2014), and the nascent SecM chain is known to interact with the negatively charged surface of the outer ribosome to stabilize the arrest (Muta et al., 2020). The arrest by SecM AP is thought to be susceptible to a pulling force exerted by the nascent chain outside the ribosome (Goldman et al., 2015). Simulations of the stalling by SecM AP showed that the structure of the nascent peptide tens of angstrom distal from the active catalytic center of the ribosome and its interaction with the exit tunnel are important for ribosome arrest (Zimmer et al., 2021). On the basis of the proposed mechanism of SecM AP and our results combining SecM AP and MSKIK on translation efficiency, we postulate that the nascent chain MSKIK, which presents just before SecM AP within the ribosomal exit tunnel, alters the native interaction between SecM AP and the ribosome to prevent or disrupt the arrest. Understanding how the nascent MSKIK peptide causes structural changes to SecM AP or acts directly on the movement of the ribosome is of great interest. The distance between SKIK and AP appears to be an important factor because the translation enhancing effect was reduced when the distance between the SKIK and the AP was increased.

Chloramphenicol physically prevents the formation of the transition state of peptide-bond formation during translation. Chloramphenicol stalls the ribosome at a specific mRNA location, and the CmlA leader ORF is a chloramphenicol-dependent AP (Gu et al., 1994). Surprisingly, N-terminal SKIK-tagging increased protein production even in the presence of chloramphenicol (Figure 4). Inhibition of protein synthesis by chloramphenicol is most effective when A, S, and T are present in the nascent peptide inside the ribosome, whereas the inhibitory effect of the drug is counteracted by the presence of G at the C-terminus of the nascent chain or the presence of G in the incoming aminoacyl-tRNA (Marks et al., 2016). However, the details of these mechanisms remain unresolved. Based on these reports and the results herein, it is plausible that the nascent chain MSKIK promotes peptidyl bond formation that occurs directly after MSKIK is created. We hypothesize that the nascent chain prevents the subsequent arrest because the MSKIK peptide is generated before AP in the translation process. As for WPPP, no arrest cancellation was observed by SKIK-tagging it was considered to be strongly arrested, but more detailed analysis is required because the distance between SKIK and AP may also be an important factor. Leininger et al. recently demonstrated using multiscale modeling simulations that consecutive positively charged residues slow down translation by generating large forces that move the P-site amino acid away from the A-site amino acid (Leininger et al., 2021). Using such a simulation approach, we postulate that explaining the translation-enhancing mechanism may be through mechanical forces generated between MSKIK and RAPs during translation. In addition, because many RAPs other than SecM or CmlA leader have been found (Su et al., 2021),and a recent study showed a force-sensitive arrest by MifM in *Bacillus subtilis* was cancelled by N-terminally adjacent dynamic nascent chain (Fujiwara et al., 2020), we expect that there are still many translation-promoting peptides remain to be undiscovered.

In conclusion, we have discovered that the nascent chain MSKIK cancels the arrest by RAPs, such as SecM and CmlA leader. We also showed that this effect was particularly pronounced when MSKIK was placed N-terminally adjacent to SecM AP. Although our data cannot reveal whether MSKIK prevents or releases the arrest, the new insights obtained herein provide important clues to further understand the translation process by the ribosome in terms of regulation of translation by nascent chains.

## ACKNOWLEDGMENTS

This research was supported by a Grant-in-Aid for JSPS Research Fellow 17J02835 and KAKENHI Grant numbers 18K14392 and 20K05806. We thank Dr. Masataka Minami (NU Protein Corporation) and Dr. Takashi Kanamori (Gene Frontiers Inc.) for providing the transcription kit and *E. coli* in vitro translation kit PUREfrex, respectively. We thank Edanz (https://jp.edanz.com/ac) for editing a draft of this manuscript.

## AUTHOR CONTRIBUTIONS

Conceptualization, T.O and H.N.; funding acquisition, T.O.; investigation, Y.N., T. O., and Y. F.; visualization, Y.N., T.O., Y.F., and T.K.; supervision, T.O. and H.N.; writing-review & editing, Y.N., T.O., Y.F., and H.N.

## INCLUSION AND DIVERSITY STATEMENT

The authors do not have relevant information to report.

## STAR METHODS

### KEY RESOURCES TABLE

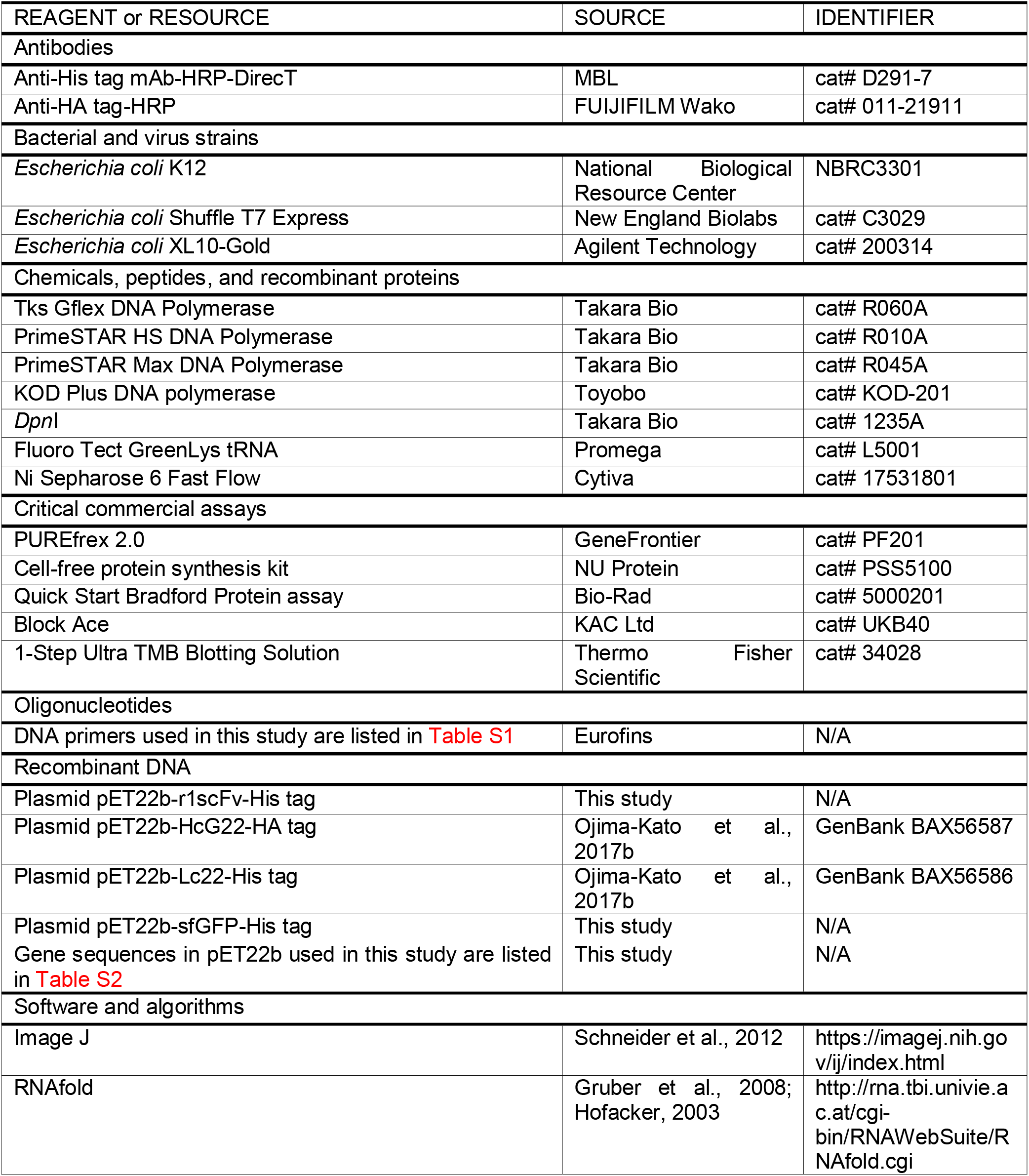

### RESOURCE AVAILABILITY

#### Lead Contact

Further information and requests for resources and reagents should be directed to and will be fulfilled by the lead contact, Teruyo Ojima-Kato.

#### Materials availability

All unique/stable reagents generated in this study are available from the lead contact with a completed materials transfer agreement.

#### Data and Code Availability

Information of DNA sequences constructed in this study is available in Table S2.

## METHOD DETAILS

### Construction of 36 SKIK codon variants

The originally used codons encoding SKIK, i.e., TCT AAA ATA AAA, were converted into 35 different patterns consisting of Ser (TCT, TCC, TCA, TCG, AGT or AGC), Lys (AAA or AAG), Ile (ATT, ATC or ATA), and Lys (AAA). The final Lys codon was fixed as AAA (Table 1). To convert the codons, we performed PCR using each primer listed in Table S1 and the plasmid pET22b-r1scFv-His tag harboring the rabbit scFv antibody gene as the template, which was challenging to express in *E. coli* (Ojima-Kato et al., 2017a, 2015). The PCR program was: 94°C 3 min; (94°C, 30 s; 55°C, 30 s; 72°C, 5 min) × 25 cycles; and 72°C, 3 min, using KOD plus (Toyobo, Osaka, Japan). The *Dpn*I (Takara Bio Inc., Otsu, Japan) treated PCR products were directly incorporated into *E. coli* XL10-Gold competent cells (Agilent Technology, Santa Clara, CA). The colonies formed on Luria–Bertani (LB) agar were checked by colony-directed PCR (94°C, 3 min; [94°C, 30 s; 55°C, 15 s; 72°C, 30 s] × 25 cycles; 72°C, 3 min; 12°C, ∞) using Tks Gflex DNA polymerase (Takara Bio Inc.) with primers F1 and R1, both of which anneal outside the T7 promoter and terminator regions (Table S1). Unless otherwise stated, 100 mg/L ampicillin was included in all cultures. The sequences of whole DNA fragments were confirmed by Sangar sequencing with the same primers. Plasmids from the selected clones were prepared using the FastGene Plasmid Mini Kit (NIPPON Genetics Co., Ltd., Tokyo, Japan), according to the manufacturer’s manual.

### Expression in *E. coli* by IPTG induction

*E. coli* SHuffle T7 Express (New England Biolabs, Ipswich, MA), which is used to express disulfide bond-containing proteins such as antibodies (Jeong et al., 2017; Robinson et al., 2015), was used as a host in this study. *E. coli* SHuffle T7 Express cells were transformed with each of the 36 plasmids that contained different codons for SKIK (Table 1). The transformants obtained on LB agar plates were inoculated into 4 mL of liquid LB medium and cultured at 37°C for 20 h as pre-cultures. A portion of these cultures (40 μL) was transferred to 4 mL of fresh LB medium and cultivated with shaking at 37°C for ∼4 h until the OD_600_ reached 0.4–0.5. Four microliters of 0.1 M IPTG were then added. Protein expression was performed at 30°C for 3 h. All cultures were grown with a shaking rate of 110 rpm. The cells were harvested by centrifugation (13,000 g, 2 min, 4°C), and the cell density was adjusted to be the same in each sample by suspending with an appropriate volume of phosphate-buffered saline (PBS; 137 mM NaCl, 8.1 mM Na_2_HPO_4_, 2.68 mM KCl, 1.47 mM KH_2_PO_4_, pH 7.4), according to the OD_600_ value after induction. Subsequently, 1 µL of *E. coli* cells suspended in approximately 200 µL PBS was mixed with a further 6 µL PBS, 1 µL 100 mM dithiothreitol (DTT), and 2 µL 5× Laemmli sample buffer (10 w/v% SDS, 50 v/v% glycerol, 30 mM Tris-HCl [pH 6.8], 0.05 w/v% Bromophenol blue) and boiled for 3 min. Without separating soluble and insoluble fractions, the samples were subjected to SDS-PAGE (12.5% acrylamide gel) and Coomassie Brilliant Blue (CBB) staining.

### Expression in *E. coli* by autoinduction

The pre-culture cells (40 μL) described above were transferred into 4 mL of Terrific Broth (TB) liquid medium (20 g Tryptone, 5 g yeast extract, 5 g NaCl, 6 g Na_2_HPO_4_, and 3 g KH_2_PO_4_ per 1 L, pH 7.2) containing 2 µL 10% glucose and 100 µL 8% lactose, and incubated at 30°C for 72 h with shaking at 110 rpm (Studier, 2005). Cell samples were harvested as described above and suspended in PBS to give equivalent OD_600_ values. Five microliters of the bacterial suspensions were subjected to SDS-PAGE and stained with CBB.

### In vitro transcription

DNA fragments containing the T7 promoter, rabbit antibody genes (HcG_22 and Lc_22 in Table S2), and T7 terminator were amplified with primer pair F1 and R1 by Tks Gflex DNA polymerase using a plasmid based on pET22b as a template. PCR products were purified by using a silica membrane column (Econospin, Ajinomono BioPharma, Ibaragi, Japan) and eluted with Tris-EDTA (TE) buffer (10 mM Tris-HCl, 1 mM EDTA-2Na, pH 8.0). *In vitro* transcription and purification of mRNA were carried out according to the cell-free protein synthesis kit instructions (NU Protein, Tokushima, Japan). In detail, the following were mixed and incubated at 37°C for 3 h: 2.5 μL 10x transcription buffer, 2.5 μL 25 mM dNTPs, 1.25 μL 0.1 M DTT, 15 μL DEPC water, 1 μL T7 RNA polymerase, and 2.5 μL DNA fragment (40–60 ng) as the template. After the reaction, 10 μL 4 M ammonium acetate and 100 μL chilled 99.5% ethanol were added sequentially. Precipitants after centrifugation (13,000 g, 20 min, 4°C) were dried under atmospheric conditions and dissolved with TE buffer.

### Protein expression in the *E. coli* in vitro expression system

The above-transcribed products (mRNA) or purified PCR products containing the T7 promoter and terminator were used as templates for the *E. coli* reconstituted cell-free protein synthesis system (PURE system) (PUREfrex® 2.0 and DS supplement, Gene Frontier Inc., Kashiwa, Japan). The reaction was performed at the 10 or 20 μL scale and 37°C for 90 min, as recommended by the manufacturer. Fluorescent Lys (fLys)-tRNA (Promega, Madison, WI, USA) was also included at 1/10 the reaction volume if required. For analysis, 5 µL of the reaction solution, 2 µL PBS, 1 µL 100 mM DTT, and 2 µL 5× sample buffer were mixed and boiled for 3 min, subjected to SDS-PAGE, and analyzed using the Typhoon FLA 9000 (Cytiva, Tokyo, Japan) laser imaging scanner to detect proteins incorporating fLys. A reaction without the template was also performed as a negative control.

### Construction of the other amino acid-encoding codons

Several peptide tags consisting of three amino acids were created and evaluated to observe possible changes with combinations of amino acids other than SKIK. Here we constructed 18 different plasmids consisting of peptide tags with SKX, KKX, and AKX (X = G [codon GGT], L [TTA], H [CAT], Y [TAT], E [GAA], and F [TTT]). The amino acid residues corresponding to X were selected to cover different properties. Insertion of tags was carried out with the same method as for constructing 36 variants using each primer pair named as three letters amino acid-F and -R, as listed in Table S1. In vivo expression following IPTG induction and whole cell SDS-PAGE analysis were performed using the same procedure described above.

### Construction of SecM containing plasmids

The SecM gene (Gene ID: 944831) was amplified from the *E. coli* K12 strain (NBRC3301) obtained from National Biological Resource Center (Kisarazu, Japan). The genome was extracted by boiling the cell pellet collected from a one-night 1 mL culture in LB medium with 0.2 mL 25 mM NaOH solution for 10 min and then neutralized by adding 32 μL 1 M Tris-HCl (pH 7.0). This genome was used as the template for the amplification of SecM with KOD plus DNA polymerase. The primers 5’-GCCGAACCAAACGCGCCCGCAAAAG-3’ and 5’-GGTGAGGCGTTGAGGGCCAGCAC-3’ were designed for cloning the full-length SecM gene except for the signal sequence (Table S1). A total of eight plasmids encoding polypeptides presented in Figure 4A were constructed by rearranging the order of the genes and connecting them using the HiFi assembly technique (New England Biolabs). The ATG start codon containing the *Nde*I site of pET22b was retained in all constructs. The DNA sequences constructed here are summarized in Table S2.

### Evaluating the combination of SKIK-CmlA leader and SKIK-WPPP

We constructed plasmids with the basic structure of AP-sfGFP-His tag with and without N-terminal SKIK to investigate the effect of SKIK peptides on non-SecM RAPs. The *E. coli* CmlA leader (KNAD), whose arrest is induced by chloramphenicol (Marks et al., 2016), and the artificially designed AP, WPPP (FQKYGIWPPP), were selected (Peil et al., 2013). The sfGFP gene was synthesized by GenScript (Pédelacq et al., 2006), Japan (Tokyo, Japan) and cloned into pET22b. In general, each plasmid was constructed by inverse PCR with primers containing the corresponding peptide-coding nucleotides and pET22b-sfGFP as the template and subsequently self-cyclized by HiFi assembly followed by transformation into *E. coli*. In the case of constructing pET22b-SecM AP-sfGFP-His tag and pET22b-SKIK-SecM AP-sfGFP-His tag, the SecM(-AP) ORFs in the constructs 5 and 6 shown in Figure 3A were replaced to sfGFP by assembling with linearized vectors. After purifying the plasmid, PCR amplified products with primer F1 and R1 were subjected to cell-free protein synthesis, as described above. For CmlA leader, the final concentration of chloramphenicol was adjusted to 0.1 to 10 μg/mL by adding 1/10 volume of 1 to 100 μg/mL chloramphenicol in ethanol into the reaction mixture.

### Protein analysis by CBB staining, dot blot, and Western blotting

After SDS-PAGE, proteins were analyzed by CBB staining and fLys detection for in vivo and in vitro expression, respectively. For scFv expressed by IPTG induction, dot blot analysis was also performed to quantify the expressed proteins in several series. In detail, the collected bacterial cells were suspended in 1 mL PBS, centrifuged (13,000 g, 2 min, 4°C), and resuspended in 3 mL 6 M Guanidine Hydrochloride (GuHCl) buffer (200 mM NaCl, 100 mM Tris-HCl, 6 M GuHCl, 10 mM DTT, pH 8.0). The cells were disrupted by sonication, then 1 μL of this sample was dropwise added to a nitrocellulose membrane and allowed to dry. Purified scFv of known concentrations were also spotted for calibration. Next, the sample was soaked in blocking buffer (1 w/v% Block Ace, KAC Ltd., Amagasaki, Japan) for 1 h. After washing with PBS-T (10 min x 3 times), the membrane was reacted with a solution of anti-His HRP-conjugate (Medical & Biological Laboratories Co., Ltd., Nagoya, Japan, 5,000x dilution with PBS) for 1 h at room temperature with gentle shaking. Then, after washing with PBS-T (as described above), 1-Step Ultra TMB Blotting Solution (Thermo Fisher Scientific K.K., Tokyo, Japan) was added to visualize the His-tagged proteins. All reactions by dot blotting were performed in six series, and the results were quantified using ImageJ software (Schneider et al., 2012) (https://imagej.nih.gov/ij/index.html). The standard His-tagged scFv used here was purified with Ni-sepharose resin in advance (0.61 mg/mL; calculated by the Bradford method). For Western blotting, 1 µL of the reaction solution after cell-free protein synthesis, 2.5 µL sterile water, 0.5 µL 0.1 M DTT, and 1 µL 5× sample buffer were mixed, boiled for 3 min, and subjected to SDS-PAGE analysis. The proteins were transferred onto a nitrocellulose membrane by a Transblot SD cell system (Bio-Rad, Hercules, CA) with standard buffer (25 mM Tris, 192 mM Glycine, 20 % MeOH, pH 8.3) at 10 V for 30 min. Subsequent operations were the same as above, but anti-HA tag-HRP (FUJIFILM Wako Pure Chemical, 10,000x dilution with PBS) was used for detecting HA-tag fused proteins.

### Fluorescence analysis

The GFP fluorescence intensity was measured as follows. Reaction solutions in the cell-free protein synthesis system were each diluted 10-fold with water following the reaction and 40 µL per well dispensed into Flat Bottom Microfluor Plates Black (Thermo Fisher Scientific), and the fluorescence intensity was measured (excitation 485 nm (bandwidth 9 nm)/emission 535 nm (bandwidth 20 nm)) using a microplate reader (Tecan Infinite 200 pro (Tecan, Männedorf, Switzerland)).

### Free energy of RNA

The minimum free energies and optimal secondary structures of mRNAs of each clone were predicted by RNAfold (http://rna.tbi.univie.ac.at/cgi-bin/RNAWebSuite/RNAfold.cgi(Gruber et al., 2008; Hofacker, 2003).

## SUPPLEMENTAL INFORMATION

Supporting data and materials are found at supplemental information

