## Supplementary material for "Nascent MSKIK peptide prevents or releases translation arrest in *Escherichia coli*": Figure S1

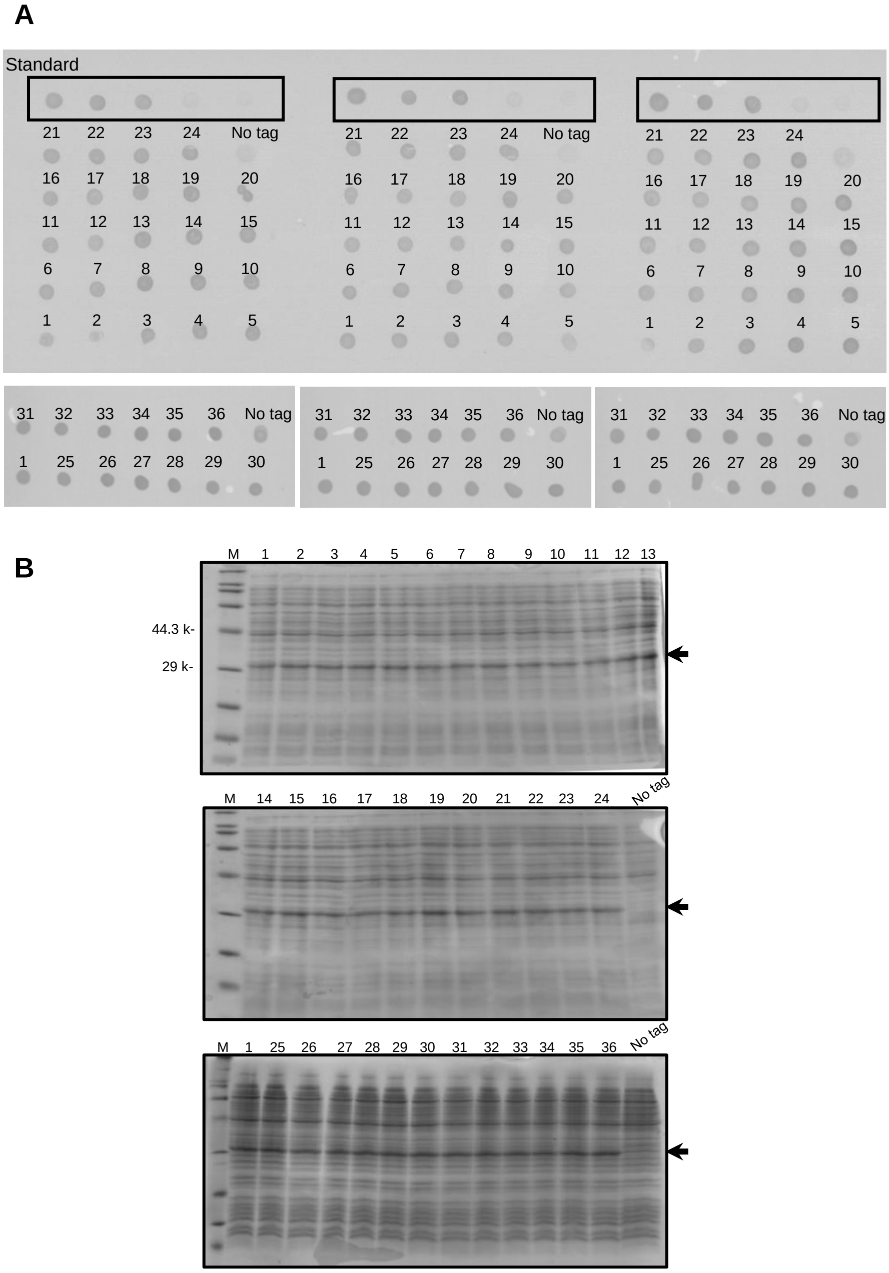


**Figure S1. Comparison of protein production by dot blot and SDS-PAGE analysis**

This data supports Figure 1 in the main manuscript. The numbers correspond to those of clones in Table 1. M: size marker.

(A) Three-fold diluted cell lysate samples and non-diluted no-tagged samples were visualized with TMB color development. The triplicated results are shown.

(B) CBB staining images of cell lysates of 36 variants and no-tagged scFvs. The arrows indicate the target protein band.
