## Supplementary material for "Nascent MSKIK peptide prevents or releases translation arrest in *Escherichia coli*": Figure S2

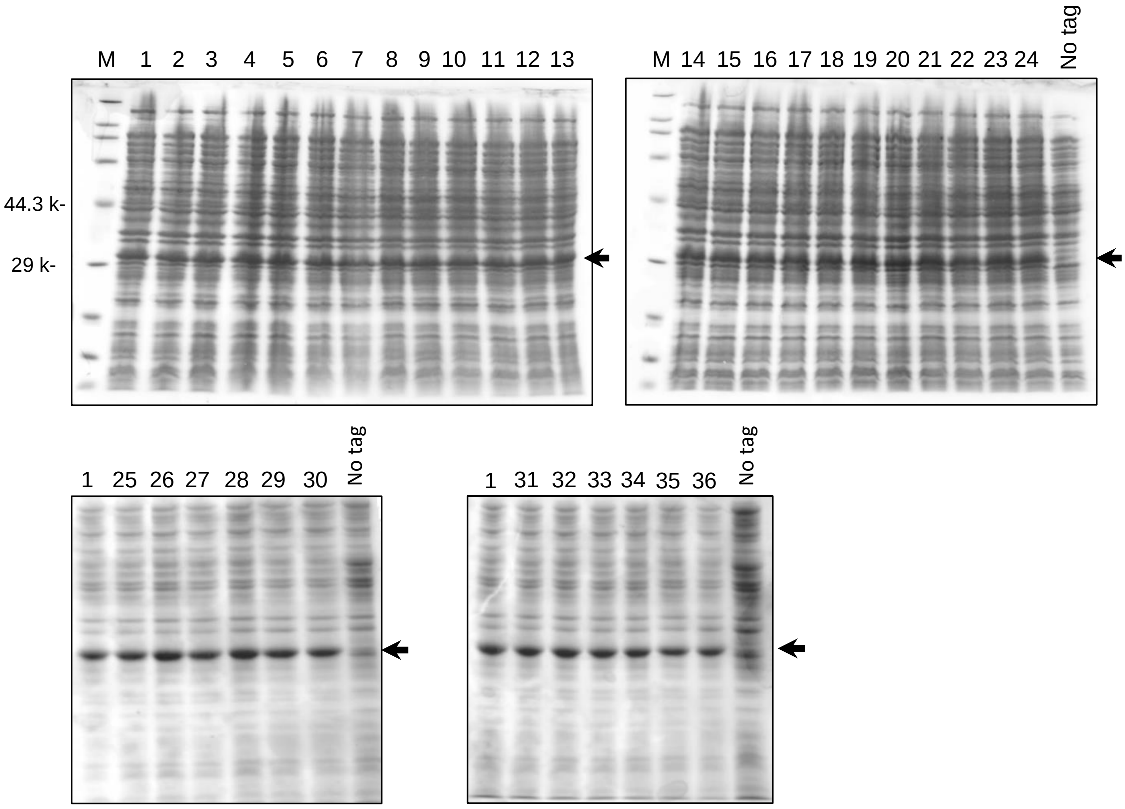


**Figure S2. SDS-PAGE of auto-induction samples**

CBB-stained SDS-PAGE gels for expression of scFv in *E. coli* Shuffle T7 Express cells by autoinduction at 30˚C for 72 h. M: size marker. The numbers correspond to the codon patterns listed in Table 1. The arrows indicate the band representing the protein of interest.
