## Supplementary material for "Nascent MSKIK peptide prevents or releases translation arrest in *Escherichia coli*": Figure S3

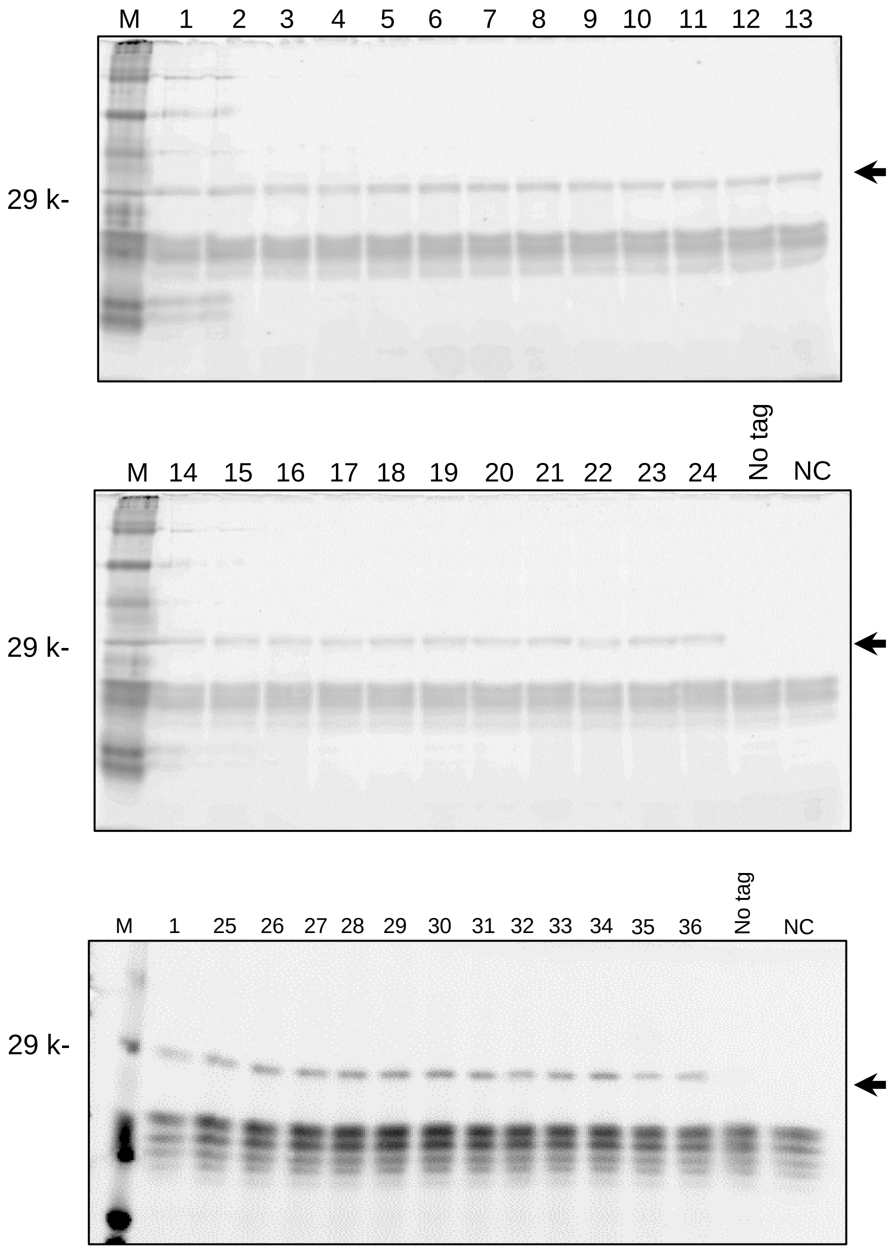


**Figure S3. Fluorescence detection of SDS-PAGE gels of cell-free protein synthesis products**

Cell-free protein synthesis products incorporating fLys t-RNA with the 36 SKIK codon variants were analyzed by fluorescence detection using a Typhoon FLA9000. M: size marker; NC: negative control. The bands indicated by the arrows are the protein of interest.
