## Supplementary material for "Nascent MSKIK peptide prevents or releases translation arrest in *Escherichia coli*": Figure S4

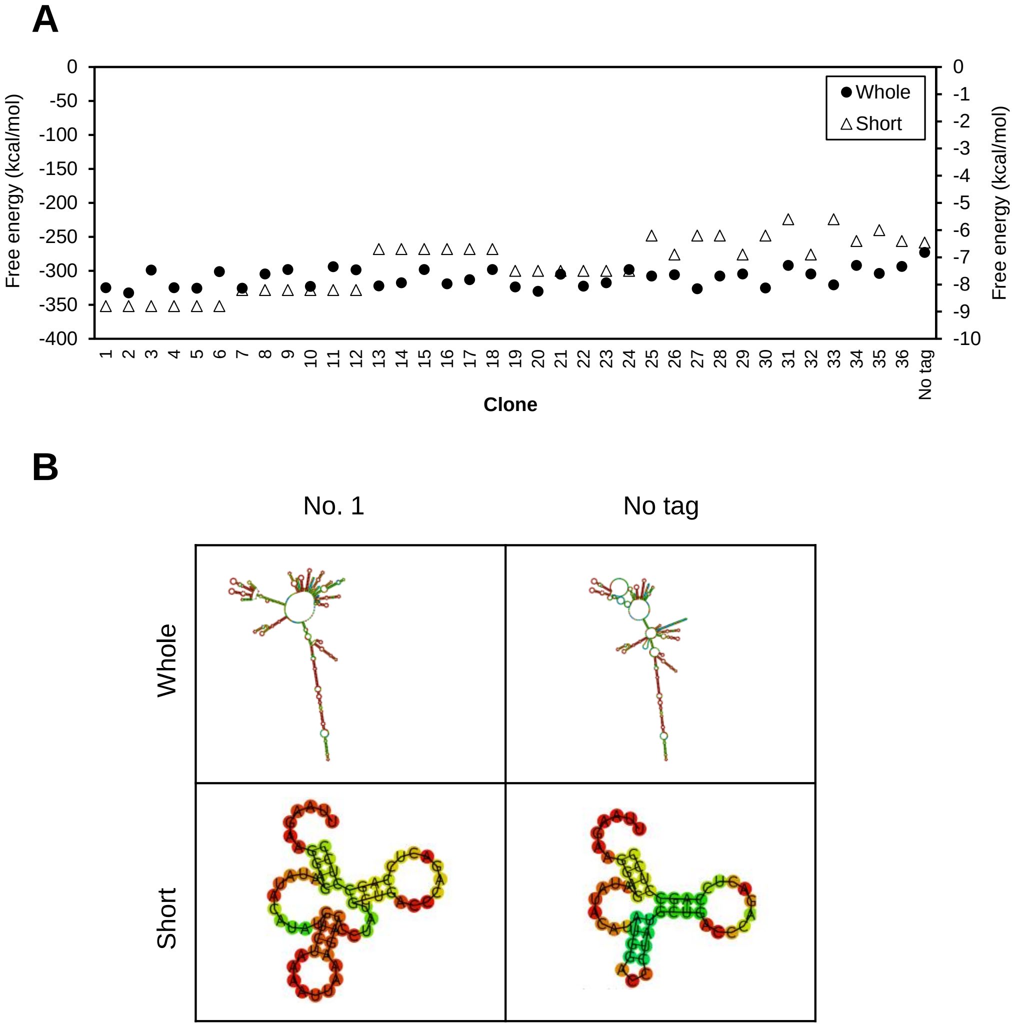


**Figure S4. The minimum free energies and optimal secondary structures of the mRNAs of the 36 SKIK variants and no-tagged mRNA**

The values and structures were predicted by RNAfold. Whole represents the entire mRNA from the transcription starting point to the T7 terminator. Short represents a focused region consisting of 64 bases that contains the Shine-Dalgano, start codon, and SKIK-encoding sequences.

(A) Minimum free energy of the centroid secondary structure in dot-bracket notation is shown for each variant. Clone numbers correspond to the SKIK-encoding codon variants listed in Table 1. The left and right y-axes show the calculated energy values for whole and short constructs, respectively.

(B) The centroid secondary structures presented correspond to SKIK-tagged clone No. 1 and No tag.
