## Supplementary material for "Nascent MSKIK peptide prevents or releases translation arrest in *Escherichia coli*": Figure S5

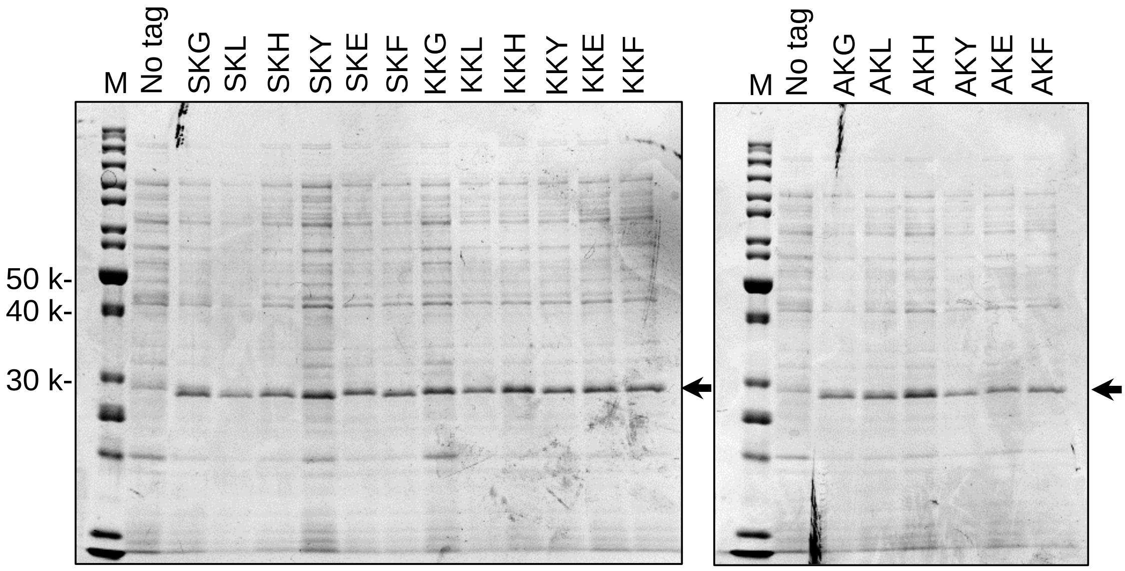


**Figure S5. Protein production in *E. coli* with N-terminal SKX, KKX, and AKX tags**

CBB staining of SDS-PAGE gels is shown. Arrows indicate bands representing the target product. M: size marker.
