## Supplementary material for "Nascent MSKIK peptide prevents or releases translation arrest in *Escherichia coli*": Table S1

Table S1. DNA Primers used in this study

| Name | Sequence (5'→3') | Discription |
|---|---|---|
| SKIK1111F | ATGTCTAAAATTAAAGACCCTATGCTGACC | |
| SKIK1111R | GGTCTTTGATTTTAGACATATGTATATCTC | |
| SKIK1121F | ATGTCTAAAATCAAAGACCCTATGCTGACC | |
| SKIK1121R | GGTCTTTGATTTTAGACATATGTATATCTC | |
| SKIK1131F | ATGTCTAAAATAAAAGACCCTATGCTGACC | |
| SKIK1131R | GGTCTTTGATTTTAGACATATGTATATCTC | |
| SKIK1211F | ATGTCTAAGATTAAAGACCCTATGCTGACC | |
| SKIK1211R | GGTCTTTAATTTTAGACATATGTATATCTC | |
| SKIK1221F | ATGTCTAAGATCAAAGACCCTATGCTGACC | |
| SKIK1221R | GGTCTTTGATCTTAGACATATGTATATCTC | |
| SKIK1231F | ATGTCTAAGATAAAAGACCCTATGCTGACC | |
| SKIK1231R | GGTCTTTTATCTTAGACATATGTATATCTC | |
| SKIK2111F | ATGTCCAAAATCTAAAGACCCTATGCTGACC | |
| SKIK2111R | GGTCTTTAATTTTGGACATATGTATATCTC | |
| SKIK2121F | ATGTCCAAAATCAAAGACCCTATGCTGACC | |
| SKIK2121R | GGTCTTTGATTTTGGACATATGTATATCTC | |
| SKIK2131F | ATGTCCAAAATAAAAGACCCTATGCTGACC | |
| SKIK2131R | GGTCTTTTATTTTGGACATATGTATATCTC | |
| SKIK2211F | ATGTCCAAGATTAAAGACCCTATGCTGACC | |
| SKIK2211R | GGTCTTTAATCTTGGACATATGTATATCTC | |
| SKIK2221F | ATGTCCAAGATCAAAGACCCTATGCTGACC | |
| SKIK2221R | GGTCTTTGATCTTGGACATATGTATATCTC | |
| SKIK2231F | ATGTCCAAGATAAAAGACCCTATGCTGACC | |
| SKIK2231R | GGTCTTTGATCTTGGACATATGTATATCTC | |
| SKIK3111F | ATGTCAAAAATTAAAGACCCTATGCTGACC | |
| SKIK3111R | GGTCTTTAATTTTTGACATATGTATATCTC | |
| SKIK3121F | ATGTCAAAAATCAAAGACCCTATGCTGACC | |
| SKIK3121R | GGTCTTTGATTTTTGACATATGTATATCTC | |
| SKIK3131F | ATGTCAAAAATAAAAGACCCTATGCTGACC | |
| SKIK3131R | GGTCTTTTATTTTAGACATATGTATATCTC | |
| SKIK3211F | ATGTCAAAGATTAAAGACCCTATGCTGACC | |
| SKIK3211R | GGTCTTTAATCTTTGACATATGTATATCTC | |
| SKIK3221F | ATGTCAAAGATCAAAGACCCTATGCTGACC | |
| SKIK3221R | GGTCTTTAATCTTTGACATATGTATATCTC | |
| SKIK3231F | ATGTCAAAGATAAAAGACCCTATGCTGACC | |
| SKIK3231R | GGTCTTTTATCTTTGACATATGTATATCTC | |
| SKIK4111F | ATGTCGAAAATTAAAGACCCTATGCTGACC | |
| SKIK4111R | GGTCTTTAATTTTCGACATATGTATATCTC | |
| SKIK4121F | ATGTCGAAAATCAAAGACCCTATGCTGACC | |
| SKIK4121R | GGTCTTTGATTTTCGACATATGTATATCTC | |
| SKIK4131F | ATGTCGAAAATAAAAGACCCTATGCTGACC | |
| SKIK4131R | GGTCTTTTATTTTCGACATATGTATATCTC | |
| SKIK4211F | ATGTCGAAGATTAAAGACCCTATGCTGACC | |

| | |
|---|---|
| SKIK4211R | GGTCTTTAATCTTCGACATATGTATATCTC |
| SKIK4221F | ATGTCGAAGATCAAAGACCCTATGCTGACC |
| SKIK4221R | GGTCTTTGATCTTCGACATATGTATATCTC |
| SKIK4231F | ATGTCGAAGATAAAAGACCCTATGCTGACC |
| SKIK4231R | GGTCTTTTATCTTCGACATATGTATATCTC |
| SKIK5111F | ATGAGTAAAATTAAAGACCCTATGCTGACC |
| SKIK5111R | GGTCTTTAATTTTACTCATATGTATATCTC |
| SKIK5121F | ATGAGTAAAATCAAAGACCCTATGCTGACC |
| SKIK5121R | GGTCTTTGATTTTACTCATATGTATATCTC |
| SKIK5131F | ATGAGTAAAATAAAAGACCCTATGCTGACC |
| SKIK5131R | GGTCTTTTATTTTACTCATATGTATATCTC |
| SKIK5211F | ATGAGTAAGATTAAAGACCCTATGCTGACC |
| SKIK5211R | GGTCTTTAATCTTACTCATATGTATATCTC |
| SKIK5221F | ATGAGTAAGATCAAAGACCCTATGCTGACC |
| SKIK5221R | GGTCTTTGATCTTACTCATATGTATATCTC |
| SKIK5231F | ATGAGTAAGATAAAAGACCCTATGCTGACC |
| SKIK5231R | GGTCTTTTATCTTACTCATATGTATATCTC |
| SKIK6111F | ATGAGCAAAATTAAAGACCCTATGCTGACC |
| SKIK6111R | GGTCTTTAATTTTGCTCATATGTATATCTC |
| SKIK6121F | ATGAGCAAAATCAAAGACCCTATGCTGACC |
| SKIK6121R | GGTCTTTGATTTTGCTCATATGTATATCTC |
| SKIK6131F | ATGAGCAAAATAAAAGACCCTATGCTGACC |
| SKIK6131R | GGTCTTTTATTTTGCTCATATGTATATCTC |
| SKIK6211F | ATGAGCAAGATTAAAGACCCTATGCTGACC |
| SKIK6211R | GGTCTTTAATCTTGCTCATATGTATATCTC |
| SKIK6221F | ATGAGCAAGATCAAAGACCCTATGCTGACC |
| SKIK6221R | GGTCTTTGATCTTGCTCATATGTATATCTC |
| SKIK6231F | ATGAGCAAGATAAAAGACCCTATGCTGACC |
| SKIK6231R | GGTCTTTTATCTTGCTCATATGTATATCTC |
| r1scFv_SKG-F | ATGTCTAAAGGTGACCCTATGCTGACC |
| r1scFv_SKG-R | GGTCACCTTTAGACATATGTATATCTC |
| r1scFv_SKL-F | ATGTCTAAATTAGACCCTATGCTGACC |
| r1scFv_SKL-R | GGTCTAATTTAGACATATGTATATCTC |
| r1scFv_SKH-F | ATGTCTAAACATGACCCTATGCTGACC |
| r1scFv_SKH-R | GGTCATGTTTAGACATATGTATATCTC |
| r1scFv_SKY-F | ATGTCTAAATATGACCCTATGCTGACC |
| r1scFv_SKY-R | GGTCATATTTAGACATATGTATATCTC |
| r1scFv_SKE-F | ATGTCTAAAGAAGACCCTATGCTGACC |
| r1scFv_SKE-R | GGTCTTCTTTAGACATATGTATATCTC |
| r1scFv_SKF-F | ATGTCTAAATTTGACCCTATGCTGACC |
| r1scFv_SKF-R | GGTCAAATTTAGACATATGTATATCTC |
| r1scFv_KKG-F | ATGAAAAAAGGTGACCCTATGCTGACC |
| r1scFv_KKG-R | GGTCACCTTTTTTCATATGTATATCTC |
| r1scFv_KKL-F | ATGAAAAAATTAGACCCTATGCTGACC |
| r1scFv_KKL-R | GGTCTAATTTTTTCATATGTATATCTC |

| | | |
|---|---|---|
| r1scFv_KKH-F | ATGAAAAAACATGACCCTATGCTGACC | |
| r1scFv_KKH-R | GGTCATGTTTTTTCATATGTATATCTC | |
| r1scFv_KKY-F | ATGAAAAAATATGACCCTATGCTGACC | |
| r1scFv_KKY-R | GGTCATATTTTTTCATATGTATATCTC | |
| r1scFv_KKE-F | ATGAAAAAAGAAGACCCTATGCTGACC | |
| r1scFv_KKE-R | GGTCTTCTTTTTTCATATGTATATCTC | |
| r1scFv_KKF-F | ATGAAAAAATTTGACCCTATGCTGACC | |
| r1scFv_KKF-R | GGTCAAATTTTTTCATATGTATATCTC | |
| r1scFv_AKG-F | ATGGCAAAAGGTGACCCTATGCTGACC | |
| r1scFv_AKG-R | GGTCACCTTTTGCCATATGTATATCTC | |
| r1scFv_AKL-F | ATGGCAAAATTAGACCCTATGCTGACC | |
| r1scFv_AKL-R | GGTCTAATTTTGCCATATGTATATCTC | |
| r1scFv_AKH-F | ATGGCAAAACATGACCCTATGCTGACC | |
| r1scFv_AKH-R | GGTCATGTTTTGCCATATGTATATCTC | |
| r1scFv_AKY-F | ATGGCAAAATATGACCCTATGCTGACC | |
| r1scFv_AKY-R | GGTCATATTTTGCCATATGTATATCTC | |
| r1scFv_AKE-F | ATGGCAAAAGAAGACCCTATGCTGACC | |
| r1scFv_AKE-R | GGTCTTCTTTTGCCATATGTATATCTC | |
| r1scFv_AKF-F | ATGGCAAAATTTGACCCTATGCTGACC | |
| r1scFv_AKF-R | GGTCAAATTTTGCCATATGTATATCTC | |
| F1 | ATCTCGATCCCGCGAAATTAATACG | |
| R1 | TCCGGATATAGTTCCTCCTTTCAG | |
| SecM-F | GCCGAACCAAACGCGCCCGCAAAAG | SecM cloning |
| SecM-R | GGTGAGGCGTTGAGGGCCAGCAC | SecM cloning |
| CmlA_sfGFP-F | AAGAATGCGATGCAAGTAAAGGTGAAGAACTGTTTAC | insertion of CmlA leader and SKIK-CmlA leader |
| CmlA_sfGFP-R | ATCGCATTCTTCATATGGATATCTCCTTCTTAAAG | insertion of CmlA leader |
| SKIK-CmlA_sfGFP-R | ATCGCATTCTTTTTTATTTTAGACATATGGATATCTCCTTCTTAAAG | insertion of SKIK-CmlA leader |
| WPPP_sfGFP-F | CGGGATTTGGCCGCCCCCTGCAAGTAAAGGTGAAGAAC | insertion of WPPP and SKIK-WPPP |
| WPPP_sfGFP-R | CCAAATCCCGTACTTCTGGAACATATGGATATCTCCTTCTTAAAG | insertion of WPPP AP |
| SKIK-WPPP_sfGFP-R | CCAAATCCCGTACTTCTGGAATTTTATTTTAGACATATGGATATCTCCTTC | insertion of SKIK-WPPP |
| His6-TAA-F | CATCACCATCACCATCATTAAAGATCCGGC | linearization of pET22b-AP–secM(-AP) and pET22b-SKIK-AP–secM(-AP) |
| SecMAP-R | TGGTTCGGCAGGGCCAGCACGGATGCC | linearization of pET22b-AP–secM(-AP) and pET22b-SKIK-AP–secM(-AP) |
| sfGFP-F | GCTGGCCCTGCCGAACCAGCAAGTAAAGGTGAAGAACTG | insertion of sfGFP into linearized vector |
| sfGFP-R | CTTTAGTGGTGGTGGTGGTGGTGCAGTTTATACAGTTCATCCATGC | insertion of sfGFP into linearized vector |
