## Supplementary material for "Nascent MSKIK peptide prevents or releases translation arrest in *Escherichia coli*": Table S2

Table S2. Plasmid DNA sequences used in this study

| Gene | DNA sequence | Discription |
|---|---|---|
| r1scFv-His tag | ATGGACCCTATGCTGACCCAGACTCCAGCCTCCGTGTCTGCAGCTGTGGGAGGCACAGTCACCATCAAGTGC<br>CAGGCCAGTGAGAACATTTACACCTCTTTAGCCTGGTATCAGCAGAAACCAGGGCACTCTCCTAAGCTCCTGA<br>TCTATTCTGCATCCACTCTGGCATCTGGGGTCGCATCGCGGTTCAAAGGCAGTGGATCTGGGACACAGTTCA<br>CTCTCACCATCAGCGGCGTGCAGTGTGATGATGCTGCCACTTACTATTGTCAATGTAGTGCTTATGGTAGGAG<br>TGGTAATTCTTTCGGCGGAGGGACCGAGGTGGTGGTCAACGGTGATCCAGTTGCACCTACTGGAGGTGGTG<br>GATCCGGCGGTGGCGGTTCTGGTGGAGGTGGATCTCAGTCGCTGGAGGAGTCCGGGGGTCGCCTGGTCAC<br>GCCTGGGACACCCCTGACACTCACCTGCACAGTCTCTGGATTCTCCCTCAGTAGTTATGCAATGAGCTGGGT<br>CCGCCAGGCTCCAGGGAAGGGGCTGGAATGGATCGGAAGTATTGGTACTGGTGGTAGCACATACTACGCGA<br>TCTGGGCGAAAGGCCGATTCACCATCTCCAAAACCTCGACCACGGTGGGTCTGAAAATCATCAGTCCGACAA<br>CCGAGGACACGGCCACCTATTTCTGTGCCCAGAATAGTTATGGTTATGTTGGTGTTAGGGAATATTTTAAGTT<br>GTGGGGCCCAGGCACCCTGGTCACCGTCGGCGGTCATCATCATCACCATCACTAA | Basic sequence without tag is shown |
| HcG_22 with/without SKIK | ATG[TCTAAAATAAAA]CAGTCGTTGGAGGAGTCCGGGGGAGGCCTGGTCCAGCCTGAGGGATCCCTGGCACT<br>CACCTGCAAAGCCTCTGGATTCACCATCAGTAGCAGCTACTACATGTGCTGGGTCCGCCAGGCTCCAGGGAA<br>GGGGCTGGAGTGGATCGGATGCATTTATGCTGGTAGTGGTGGTACATACTACGCGAGCTGGGCGAAAGGCC<br>GATTCACCATCTCCAAGTCCTCGTCGACCACGGTGACTCTGCAAATGACCAGTCTGACAGCCGCGGACACGG<br>CCACCTATTTCTGTGCGAGGGACGTTGATGTTAGTGGTTATGGTCTGGACTTGTGGGGCCCAGGCACCCTGG<br>TCACCGTCTCCTCAGGGCAACCTAAGGCTCCATCAGTCTTCCCACTGGCACCGTGTTGTGGTGATACCCCGA<br>GCAGCACCGTTACCCTGGGTTGTCTGGTTAAAGGTTATCTGCCGGAACCGGTTACCGTTACCTGGAATAGCG<br>GCACCCTGACCAATGGTGTTCGTACCTTTCCGAGCGTTCGTCAGAGCAGCGGTCTGTATAGCCTGAGCAGCG<br>TTGTTAGCGTTACCAGCAGCAGCCAGCCGGTTACCTGTAATGTTGCACATCCGGCTACCAATACCAAAGTTGA<br>TAAAACCGTTGCACCGAGCACCTGTGGCGGTGGTGGGAGCGCCCAGCTCGAAAAGGAGCTGCAAGCCCTGG | VH-CH1 region of rabbit antibody (BAX56587) with leucine zipper and HA tag |

| | | |
|---|---|---|
| | AGAAGGAGAACGCCCAGCTCGAATGGGAGCTCCAGGCCCTGGAGAAGGAGCTGGCCCAGAAGGGCGGTACCATGTACCCATACGATGTTCCAGATTACGCTTAATAA | |
| Lc_22 with/without SKIK | ATG[TCTAAAATAAAA]GATGTTGTGATGACCCAGACTCCAGACTCCGTGTCTGCAGCTGTGGGAGGCACAGTCACCATCAATTGCCAGGCCAGTGAGAGCATTTATAGCAATTTAGCCTGGTATCAGCAGAAACCAGGGCAGCCTCCCAAGCTCCTGATCTATGCTGCATCGAAACTGGCATCTGGGGTCCCATCGCGGTTCAAAGGCAGTGGATCTGGGACACAGTTCACTCTCACCATCAGCGACCTGGAGTGTGCCGATGCTGCCACTTACTACTGTCAATGTACTTATTATGGTAGTAGTGCTGTTCCTAATGCTTTCGGCGGAGGGACCGAGGTGGTGGTCAAAGGTGATCCAGTTGCACCTACTGTCCTCATCTTCCCACCAGCTGCTGATCAGGTGGCAACTGAAACAGTCACCATCGTGTGTGTTGCCAATAAATACTTTCCGGATGTTACCGTTACCTGGGAAGTTGATGGCACCACCCAGACCACCGGTATTGAAAATAGCAAAACACCGCAGAATAGCGCAGATTGTACCTATAATCTGAGCAGCACCCTGACCCTGACCAGCACCCAGTATAACAGCCATAAAGAATATACCTGCAAAGTGACCCAGGGTACAACCAGCGTTGTTCAGAGCTTTAATCGTGGTGATTGTGGCGGTGGTGGGAGCGCCCAGCTCAAGAAGAAGCTGCAAGCCCTGAAGAAGAAGAACGCCCAGCTCAAGTGGAAGCTCCAGGCCCTGAAGAAGAAGCTGGCCCAGAAGGGCGGTTCCCATCATCATCACCATCACTAATAA | VL-CL region of rabbit antibody (BAX56586) with leucine zipper and His tag |
| secM(-AP)–AP | ATGGCCGAACCAAACGCGCCCGCAAAAGCGACAACCCGCAACCACGAGCCTTCAGCCAAAGTTAACTTTGGTCAATTGGCCTTGCTGGAAGCGAACACACGCCGCCCGAATTCGAACTATTCCGTTGATTACTGGCATCAACATGCCATTCGCACGGTAATCCGTCATCTTTCTTTCGCAATGGCACCGCAAACACTGCCCGTTGCTGAAGAATCTTTGCCTCTTCAGGCGCAACATCTTGCATTACTGGATACGCTCAGCGCGCTGCTGACCCAGGAAGGCACGCCGTCTGAAAAGGGTTATCGCATTGATTATGCGCATTTTACCCCACAAGCAAAATTCAGCACGCCCGTCTGGATAAGCCAGGCGCAAGGCATCCGTGCTGGCCCTCACATCACCATCACCATCATTAA | |

| | |
|---|---|
| SKIK–secM(-AP)–AP | ATGTCTAAAATAAAAGCCGAACCAAACGCGCCCGCAAAAGCGACAACCCGCAACCACGAGCCTTCAGCCAAA<br>GTTAACTTTGGTCAATTGGCCTTGCTGGAAGCGAACACACGCCGCCCGAATTCGAACTATTCCGTTGATTACT<br>GGCATCAACATGCCATTCGCACGGTAATCCGTCATCTTTCTTTCGCAATGGCACCGCAAACACTGCCCGTTGC<br>TGAAGAATCTTTGCCTCTTCAGGCGCAACATCTTGCATTACTGGATACGCTCAGCGCGCTGCTGACCCAGGAA<br>GGCACGCCGTCTGAAAAGGGTTATCGCATTGATTATGCGCATTTTACCCCACAAGCAAAATTCAGCACGCCCG<br>TCTGGATAAGCCAGGCGCAAGGCATCCGTGCTGGCCCTCACATCACCATCACCATCATTAA |
| secM(-AP) | ATGGCCGAACCAAACGCGCCCGCAAAAGCGACAACCCGCAACCACGAGCCTTCAGCCAAAGTTAACTTTGGT<br>CAATTGGCCTTGCTGGAAGCGAACACACGCCGCCCGAATTCGAACTATTCCGTTGATTACTGGCATCAACATG<br>CCATTCGCACGGTAATCCGTCATCTTTCTTTCGCAATGGCACCGCAAACACTGCCCGTTGCTGAAGAATCTTT<br>GCCTCTTCAGGCGCAACATCTTGCATTACTGGATACGCTCAGCGCGCTGCTGACCCAGGAAGGCACGCCGTC<br>TGAAAAGGGTTATCGCATTGATTATGCGCATTTTACCCCACAAGCAAAACATCACCATCACCATCATTAA |
| SKIK–secM(-AP) | ATGTCTAAAATAAAAGCCGAACCAAACGCGCCCGCAAAAGCGACAACCCGCAACCACGAGCCTTCAGCCAAA<br>GTTAACTTTGGTCAATTGGCCTTGCTGGAAGCGAACACACGCCGCCCGAATTCGAACTATTCCGTTGATTACT<br>GGCATCAACATGCCATTCGCACGGTAATCCGTCATCTTTCTTTCGCAATGGCACCGCAAACACTGCCCGTTGC<br>TGAAGAATCTTTGCCTCTTCAGGCGCAACATCTTGCATTACTGGATACGCTCAGCGCGCTGCTGACCCAGGAA<br>GGCACGCCGTCTGAAAAGGGTTATCGCATTGATTATGCGCATTTTACCCCACAAGCAAAACATCACCATCACC<br>ATCATTAA |
| AP–secM(-AP) | ATGTTCAGCACGCCCGTCTGGATAAGCCAGGCGCAAGGCATCCGTGCTGGCCCTGCCGAACCAAACGCGCC<br>CGCAAAAGCGACAACCCGCAACCACGAGCCTTCAGCCAAAGTTAACTTTGGTCAATTGGCCTTGCTGGAAGC<br>GAACACACGCCGCCCGAATTCGAACTATTCCGTTGATTACTGGCATCAACATGCCATTCGCACGGTAATCCGT<br>CATCTTTCTTTCGCAATGGCACCGCAAACACTGCCCGTTGCTGAAGAATCTTTGCCTCTTCAGGCGCAACATC<br>TTGCATTACTGGATACGCTCAGCGCGCTGCTGACCCAGGAAGGCACGCCGTCTGAAAAGGGTTATCGCATTG<br>ATTATGCGCATTTTACCCCACAAGCAAAACATCACCATCACCATCATTAA |

| | |
|---|---|
| SKIK–AP–secM(-AP) | ATGTCTAAAATAAAATTCAGCACGCCCGTCTGGATAAGCCAGGCGCAAGGCATCCGTGCTGGCCCTGCCGAA<br>CCAAACGCGCCCGCAAAAGCGACAACCCGCAACCACGAGCCTTCAGCCAAAGTTAACTTTGGTCAATTGGCC<br>TTGCTGGAAGCGAACACACGCCGCCCGAATTCGAACTATTCCGTTGATTACTGGCATCAACATGCCATTCGCA<br>CGGTAATCCGTCATCTTTCTTTCGCAATGGCACCGCAAACACTGCCCGTTGCTGAAGAATCTTTGCCTCTTCA<br>GGCGCAACATCTTGCATTACTGGATACGCTCAGCGCGCTGCTGACCCAGGAAGGCACGCCGTCTGAAAAGG<br>GTTATCGCATTGATTATGCGCATTTTACCCCACAAGCAAAACATCACCATCACCATCATTAA |
| secM(-AP)–SKIK–AP | ATGGCCGAACCAAACGCGCCCGCAAAAGCGACAACCCGCAACCACGAGCCTTCAGCCAAAGTTAACTTTGGT<br>CAATTGGCCTTGCTGGAAGCGAACACACGCCGCCCGAATTCGAACTATTCCGTTGATTACTGGCATCAACATG<br>CCATTCGCACGGTAATCCGTCATCTTTCTTTCGCAATGGCACCGCAAACACTGCCCGTTGCTGAAGAATCTTT<br>GCCTCTTCAGGCGCAACATCTTGCATTACTGGATACGCTCAGCGCGCTGCTGACCCAGGAAGGCACGCCGTC<br>TGAAAAGGGTTATCGCATTGATTATGCGCATTTTACCCCACAAGCAAAATCTAAAATAAAATTCAGCACGCCCG<br>TCTGGATAAGCCAGGCGCAAGGCATCCGTGCTGGCCCTCAACGCCTCACCCATCACCATCACCATCATTAA |
| secM(-AP)–MSKIK–AP | ATGGCCGAACCAAACGCGCCCGCAAAAGCGACAACCCGCAACCACGAGCCTTCAGCCAAAGTTAACTTTGGT<br>CAATTGGCCTTGCTGGAAGCGAACACACGCCGCCCGAATTCGAACTATTCCGTTGATTACTGGCATCAACATG<br>CCATTCGCACGGTAATCCGTCATCTTTCTTTCGCAATGGCACCGCAAACACTGCCCGTTGCTGAAGAATCTTT<br>GCCTCTTCAGGCGCAACATCTTGCATTACTGGATACGCTCAGCGCGCTGCTGACCCAGGAAGGCACGCCGTC<br>TGAAAAGGGTTATCGCATTGATTATGCGCATTTTACCCCACAAGCAAAAATGTCTAAAATAAAATTCAGCACGC<br>CCGTCTGGATAAGCCAGGCGCAAGGCATCCGTGCTGGCCCTCAACGCCTCACCCATCACCATCACCATCATT<br>AA |

| | |
|---|---|
| sfGFP | ATGGCAAGTAAAGGTGAAGAACTGTTTACCGGCGTGGTTCCGATTCTGGTGGAACTGGATGGTGATGTTAATG<br>GCCATAAATTCAGCGTGCGCGGCGAAGGTGAAGGCGATGCCACCAACGGTAAACTGACGCTGAAATTTATCT<br>GCACCACGGGTAAACTGCCGGTGCCGTGGCCGACCCTGGTTACCACGCTGACGTATGGCGTGCAGTGTTTC<br>AGCCGTTACCCGGATCATATGAAACGCCACGATTTCTTTAAAAGCGCCATGCCGGAAGGTTATGTTCAGGAAC<br>GTACCATTTCTTTTAAAGATGATGGCACCTACAAAACGCGCGCAGAAGTGAAATTCGAAGGTGATACCCTGGT<br>TAACCGTATTGAACTGAAAGGCATCGATTTCAAAGAAGATGGTAACATCCTGGGCCATAAACTGGAATACAAC<br>TTCAACTCTCACAACGTGTACATCACCGCAGATAAACAGAAAAACGGTATCAAAGCGAACTTCAAAATCCGCC<br>ATAATGTGGAAGATGGCAGCGTTCAGCTGGCGGATCACTATCAGCAGAACACCCCGATTGGTGATGGCCCGG<br>TTCTGCTGCCGGATAATCATTACCTGAGCACGCAGTCTGTGCTGAGTAAAGATCCGAACGAAAAACGTGATCA<br>TATGGTGCTGCTGGAATTTGTTACCGCGGCCGGTATCACGCACGGCATGGATGAACTGTATAAACTGCATCAC<br>CATCACCATCATTAA |
| SecM AP- sfGFP with/witout SKIK | ATG[TCTAAAATAAAA]TTCAGCACGCCCGTCTGGATAAGCCAGGCGCAAGGCATCCGTGCTGGCCCTGCCGA<br>ACCAGCAAGTAAAGGTGAAGAACTGTTTACCGGCGTGGTTCCGATTCTGGTGGAACTGGATGGTGATGTTAAT<br>GGCCATAAATTCAGCGTGCGCGGCGAAGGTGAAGGCGATGCCACCAACGGTAAACTGACGCTGAAATTTATC<br>TGCACCACGGGTAAACTGCCGGTGCCGTGGCCGACCCTGGTTACCACGCTGACGTATGGCGTGCAGTGTTT<br>CAGCCGTTACCCGGATCATATGAAACGCCACGATTTCTTTAAAAGCGCCATGCCGGAAGGTTATGTTCAGGAA<br>CGTACCATTTCTTTTAAAGATGATGGCACCTACAAAACGCGCGCAGAAGTGAAATTCGAAGGTGATACCCTGG<br>TTAACCGTATTGAACTGAAAGGCATCGATTTCAAAGAAGATGGTAACATCCTGGGCCATAAACTGGAATACAA<br>CTTCAACTCTCACAACGTGTACATCACCGCAGATAAACAGAAAAACGGTATCAAAGCGAACTTCAAAATCCGC<br>CATAATGTGGAAGATGGCAGCGTTCAGCTGGCGGATCACTATCAGCAGAACACCCCGATTGGTGATGGCCCG<br>GTTCTGCTGCCGGATAATCATTACCTGAGCACGCAGTCTGTGCTGAGTAAAGATCCGAACGAAAAACGTGATC<br>ATATGGTGCTGCTGGAATTTGTTACCGCGGCCGGTATCACGCACGGCATGGATGAACTGTATAAACTGCATCA<br>CCATCACCATCATTAA |

| | |
|---|---|
| CmlA leader-sfGFP with/witout SKIK | ATG[TCTAAAATAAAA]AAGAATGCGGATGCAAGTAAAGGTGAAGAACTGTTTACCGGCGTGGTTCCGATTCTG<br>GTGGAACTGGATGGTGATGTTAATGGCCATAAATTCAGCGTGCGCGGCGAAGGTGAAGGCGATGCCACCAAC<br>GGTAAACTGACGCTGAAATTTATCTGCACCACGGGTAAACTGCCGGTGCCGTGGCCGACCCTGGTTACCACG<br>CTGACGTATGGCGTGCAGTGTTTCAGCCGTTACCCGGATCATATGAAACGCCACGATTTCTTTAAAAGCGCCA<br>TGCCGGAAGGTTATGTTCAGGAACGTACCATTTCTTTTAAAGATGATGGCACCTACAAAACGCGCGCAGAAGT<br>GAAATTCGAAGGTGATACCCTGGTTAACCGTATTGAACTGAAAGGCATCGATTTCAAAGAAGATGGTAACATC<br>CTGGGCCATAAACTGGAATACAACTTCAACTCTCACAACGTGTACATCACCGCAGATAAACAGAAAAACGGTA<br>TCAAAGCGAACTTCAAAATCCGCCATAATGTGGAAGATGGCAGCGTTCAGCTGGCGGATCACTATCAGCAGA<br>ACACCCCGATTGGTGATGGCCCGGTTCTGCTGCCGGATAATCATTACCTGAGCACGCAGTCTGTGCTGAGTA<br>AAGATCCGAACGAAAAACGTGATCATATGGTGCTGCTGGAATTTGTTACCGCGGCCGGTATCACGCACGGCA<br>TGGATGAACTGTATAAACTGCATCACCATCACCATCATTAA |
| WPPP-sfGFP with/witout SKIK | ATG[TCTAAAATAAAA]TTCCAGAAGTACGGGATTTGGCCGCCCCCTGCAAGTAAAGGTGAAGAACTGTTTACC<br>GGCGTGGTTCCGATTCTGGTGGAACTGGATGGTGATGTTAATGGCCATAAATTCAGCGTGCGCGGCGAAGGT<br>GAAGGCGATGCCACCAACGGTAAACTGACGCTGAAATTTATCTGCACCACGGGTAAACTGCCGGTGCCGTGG<br>CCGACCCTGGTTACCACGCTGACGTATGGCGTGCAGTGTTTCAGCCGTTACCCGGATCATATGAAACGCCAC<br>GATTTCTTTAAAAGCGCCATGCCGGAAGGTTATGTTCAGGAACGTACCATTTCTTTTAAAGATGATGGCACCTA<br>CAAAACGCGCGCAGAAGTGAAATTCGAAGGTGATACCCTGGTTAACCGTATTGAACTGAAAGGCATCGATTTC<br>AAAGAAGATGGTAACATCCTGGGCCATAAACTGGAATACAACTTCAACTCTCACAACGTGTACATCACCGCAG<br>ATAAACAGAAAAACGGTATCAAAGCGAACTTCAAAATCCGCCATAATGTGGAAGATGGCAGCGTTCAGCTGGC<br>GGATCACTATCAGCAGAACACCCCGATTGGTGATGGCCCGGTTCTGCTGCCGGATAATCATTACCTGAGCAC<br>GCAGTCTGTGCTGAGTAAAGATCCGAACGAAAAACGTGATCATATGGTGCTGCTGGAATTTGTTACCGCGGC<br>CGGTATCACGCACGGCATGGATGAACTGTATAAACTGCATCACCATCACCATCATTAA |

The coding regions cloned into NdeI site of pET22b are shown. SKIK tag and His tag are shown in red and light blue, respectively.
